# Diurnal Regulation of Flagellar Length and Swimming Speed in the Red-Tide Raphidophyte *Chattonella marina*

**DOI:** 10.64898/2026.02.18.706705

**Authors:** Yusaku Fujita, Azusa Kage, Takayuki Nishizaka

**Affiliations:** Department of Physics, Gakushuin University, Toshima-ku, Tokyo, Japan; Graduate School of Engineering, Muroran Institute of Technology, Muroran, Hokkaido, Japan

## Abstract

The raphidophyte *Chattonella marina* is a harmful algal bloom (HAB) species known for its distinct diurnal vertical migration (DVM), a behavior important for its survival and bloom formation. However, the single-cell mechanisms governing this migration remain unclear. In this study, we investigated the swimming characteristics of individual *C. marina* cells during day (light) and night (dark) phases. We observed a strong positive correlation between the length of the propulsive anterior flagellum and the cell’s swimming speed. We discovered that the length distribution of the anterior flagellum is different during the day and at night. We also found that the beat frequency of the anterior flagellum was significantly higher during the day compared to the night. This resulted in faster mean swimming speeds during the light phase. To investigate the mechanism of length regulation, we tested the role of intraflagellar transport (IFT) using the IFT dynein inhibitor, ciliobrevin D. Treatment with ciliobrevin D induced a time- and concentration-dependent shortening of the anterior flagellum. This is the first pharmacological evidence to suggest that an IFT-like mechanism may actively control motile flagellar length in *C. marina*. These findings suggest that *C. marina* modulates its swimming speed through diurnal changes in both flagellar length and beat frequency, likely as an energy-saving strategy coupled to its DVM.

## Introduction

Harmful algal blooms (HABs), or red tides, pose a significant threat to marine ecosystems and aquaculture industries worldwide. The raphidophyte genus *Chattonella* is notorious for forming massive blooms that have caused catastrophic fish kills (Imai et al., 2006). A key ecological strategy for *Chattonella* is its pronounced diurnal vertical migration (DVM) (Shikata et al., 2023, 2015). During the day, this photosynthetic flagellate migrates upwards into the surface layer to access sunlight. At night, they descend to deeper, nutrient-rich waters. This behavior allows them to optimize both photosynthesis and nutrient uptake, and it is considered a critical factor in the initiation and maintenance of dense blooms (Watanabe et al., 1991, 1995).

While the ecological significance of DVM is well-established, the underlying physiological and mechanical changes at the single-cell level remain largely unexplored. It is unclear how individual cells modulate their swimming apparatus to reverse their migration direction or whether their intrinsic swimming properties, such as speed, change between day and night.

*Chattonella* is a unicellular biflagellate that belongs to raphidophyte. Propulsion is generated primarily by the anterior flagellum, which is covered in hair-like mastigonemes and pulls the cell forward (Hara et al., 1985; Vesk & Moestrup, 1987). A second, smooth posterior flagellum trails the cell body. The regulation of this flagellar system is central to the cell’s motility and migratory behavior.

This study aims to elucidate the single-cell basis of swimming behavior in *Chattonella marina* (Fig. 1, Supplementary Movie S1). We focused on quantifying differences in key swimming parameters, specifically flagellar length, beat frequency, and swimming speed, between the day and night phases. Furthermore, we investigated a potential molecular mechanism for flagellar length regulation by applying ciliobrevin D (Firestone et al., 2012), a known inhibitor of intraflagellar transport (IFT) dynein. Our findings provide the first evidence linking diurnal changes in flagellar length and beat frequency to the swimming behavior of *C. marina*, offering a single-cell mechanical basis for its DVM. Furthermore, we provide novel insight into the molecular regulation of flagellar length in motile flagellates, suggesting a conserved role for IFT-related machinery.

**Figure 1.**
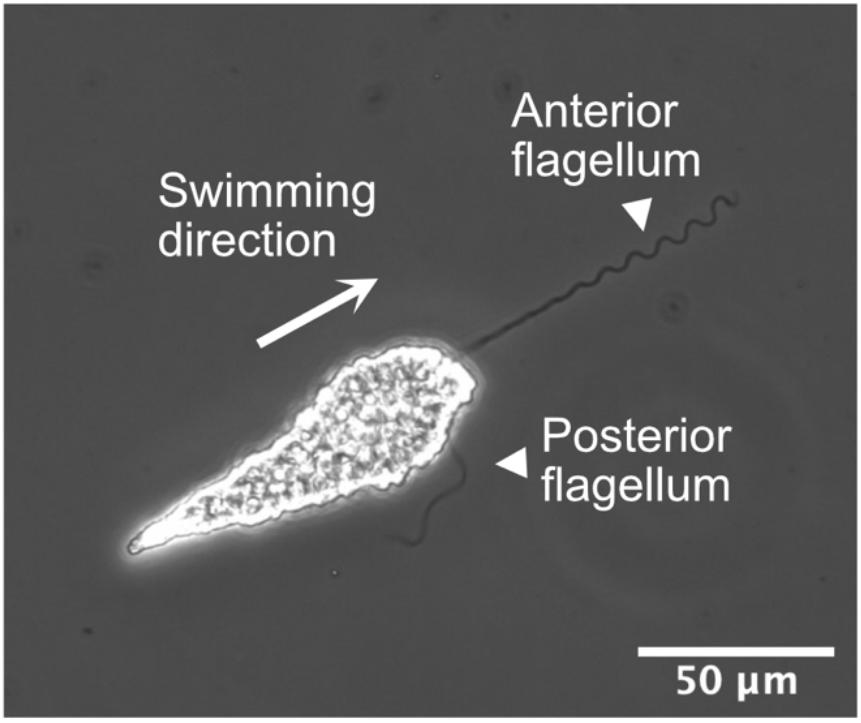
Phase-contrast micrograph of *Chattonella marina* (NIES-1 strain). The anterior (pulling) and posterior (trailing) flagella are indicated. See Supplementary Movie S1.

## Methods

### Cell Culture

*Chattonella marina* (NIES-1 strain) was cultured in SWM-3 liquid medium (Shikata et al., 2011). Cultures were maintained in an incubator (BIOTRON, Nippon Medical and Chemical Instruments, Osaka, Japan) at 25°C under a 12:12 hour light:dark cycle (light period 9:00–21:00, ca. 4000 lux; fluorescent lamp). Cells were subcultured every two weeks, and experiments were performed using cultures 7 to 10 days post-inoculation.

### Observation Chamber

A custom observation chamber was constructed from a 24×36 mm cover glass (Matsunami, Osaka, Japan), a 5 mm thick silicone rubber sheet (Kokugo, Tokyo, Japan) with a 10×10 mm central cutout, and an 18×18 mm cover glass lid (Matsunami, Osaka, Japan). 500 µL of cell culture was loaded into this chamber (Fig. 2). For detailed observation of the posterior flagellum, a spacer-less chamber was made by placing 50 µL of culture between two cover glasses (24×36 mm and 18×18 mm) and sealing the edges with vaseline to prevent evaporation and flow.

**Figure 2.**
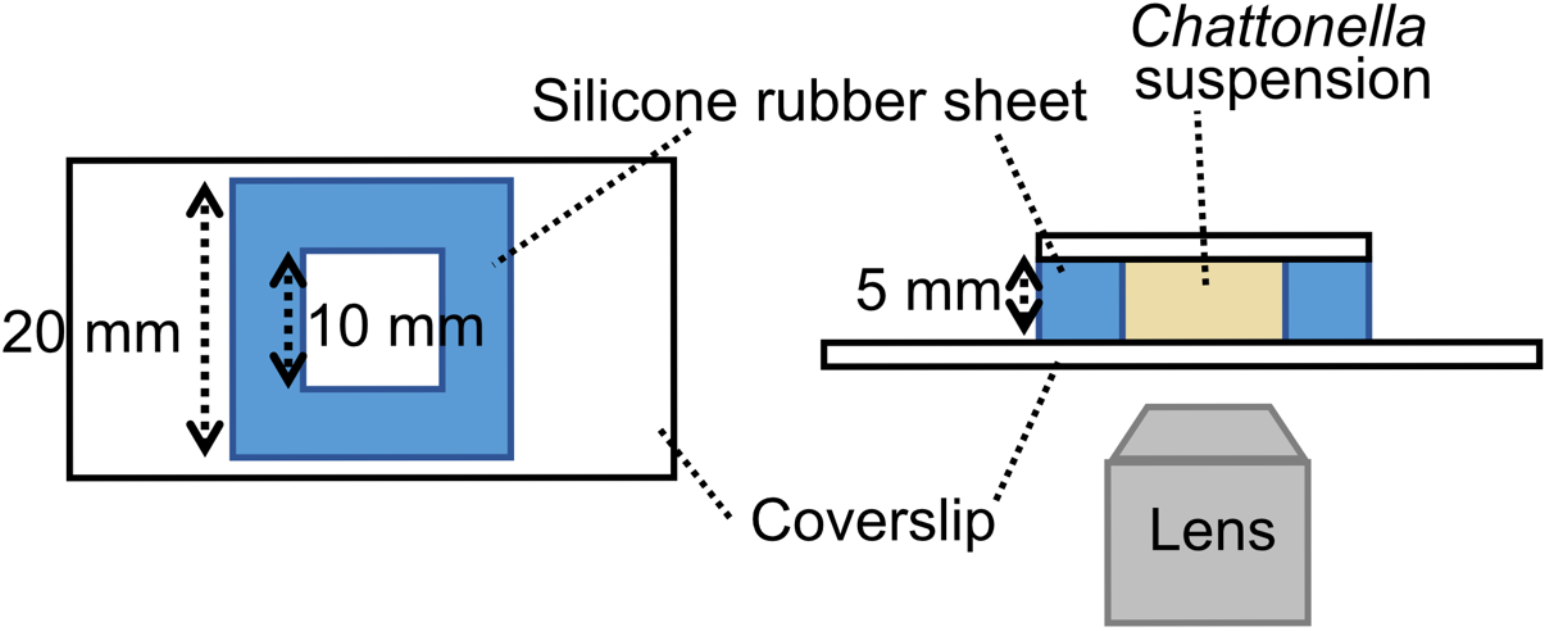
Schematic of the custom-built observation chamber used for microscopy.

### Microscopy and Image Acquisition

Observations were performed using an Olympus IX71 inverted microscope equipped with 10x (Plan 10x / 0.25) and 20x (LUCPlan FLN 20x / 0.45 Ph1) objectives and a halogen lamp (IX-HLSH100, Olympus, Tokyo, Japan) as a light source. Phase-contrast illumination was used to visualize flagella. Images were captured with a high-speed camera (VCC-H1540M, Digimo, Osaka, Japan) using Endless HDR software. Two recording conditions were used:

1. Swimming and Length Analysis: 10x objective, 30 fps, 1/500 s exposure time, 10-second duration.
2. Waveform and Frequency Analysis: 20x objective, 200 fps (for anterior flagellum) or 500 fps (for posterior flagellum), 1/500 s exposure time, 10-second duration.

### Day/Night Sampling

For diurnal comparisons, cells were sampled from the incubator during the day (10:00– 11:00) and night (22:00–23:00). To minimize mechanical stress, a P-1000 micropipette with the tip cut to widen the orifice was used for sample collection. Samples were loaded into the 5 mm deep chamber (Fig. 2) and allowed to adapt for 5-10 minutes before recording. Cells were recorded at both the top (near the 18×18 mm coverslip) and bottom (near the 24×36 mm coverslip) focal planes.

### Image and Data Analysis

#### Flagellar Length, Cell Size, and Swimming Speed

Analysis was performed using ImageJ. Flagellar length was measured as the straight-line distance from the base to the tip in frames where the flagellum was in focus. Cell length (long axis) and width (short axis) were measured similarly. Cell volume (V) was approximated as an ellipsoid: V=(4/3)×π×(long axis/2)×(short axis/2)^2^. Swimming speed was determined using the Manual Tracking plugin by tracking the cell body’s centroid over approximately a hundred frames.

#### Flagellar Waveform and Frequency

Analysis was conducted using the Bohboh software (Bohbohsoft, Tokyo, Japan) (Shiba et al., 2002). Flagella were semi-automatically traced for approximately 20 frames for each data. For frequency analysis, the curvature at a set point (s_0) along the flagellum was calculated over time. The resulting time-series data (.bol file) was exported to the regression software Bohboh_L (Bohbohsoft, Tokyo, Japan), and a sine function (Y=A0×sin(2×π×(X+A2)/A1)+A3) was fitted using the least-squares method, where X is the distance along the flagellum. The frequency was derived from the ‘A1’ parameter (period) and the frame rate.

#### Inhibitor Experiment

A 50 mM stock solution of ciliobrevin D in DMSO was thawed and diluted in SWM-3 medium to a 100 µM working stock. A 500-fold dilution of DMSO in SWM-3 was prepared as a vehicle control. *Chattonella* cultures were aliquoted (900-950 µL) into 1.5 mL tubes and treated to final concentrations of 5 µM ciliobrevin D, 10 µM ciliobrevin D, or an equivalent volume of the DMSO vehicle control. The concentrations were determined based on the preliminary experiments. A no-addition culture served as the absolute control. Cells were sampled from these tubes, loaded into observation chambers, and recorded at 0-, 3-, and 6-hours post-treatment.

#### Statistical Analysis

Statistical tests, including the Kolmogorov-Smirnov test, Mann-Whitney U-test, and Fisher’s Z-transform for comparing correlation coefficients, were performed using R or Python (scipy).

## Results

### Diurnal Changes in Swimming Behavior and Flagellar Length

We first observed the distribution of cells in the observation chamber of the depth 5 mm. During the day, a majority of cells accumulated at the top surface. At night, cells tended to accumulate at the bottom surface.

We then measured the anterior flagellar length of cells in both phases (Fig. 3). During the day, the anterior flagella were much longer (85% on average) at the top surface than at the bottom surface (top: 76.3±14.1 µm, n=45, bottom: 41.2±20.0 µm, n=26, mean±standard deviation; Mann-Whitney U-test, p=2.2×10^−9^). At night, the difference at the top and at the bottom was less marked, i.e., top was 13% longer on average, though still significant (top: 64.2±9.2 µm, n=45, bottom: 56.6±14.9 µm, n=53; Mann-Whitney U-test, p=0.02). A pooled comparison of all cells showed that the length of anterior flagella during the day was significantly different from that at night (day: n=71, night: n=98, Kolmogorov-Smirnov test, p=5.6×10^−4^). The distribution was broader during the day (mean±SD: 63.4±23.6 µm, the difference between the maximum and the minimum values: 96.7 µm) than at night (mean±SD: 60.1±13.1 µm, the difference between the maximum and the minimum values: 61.7 µm).

**Figure 3.**
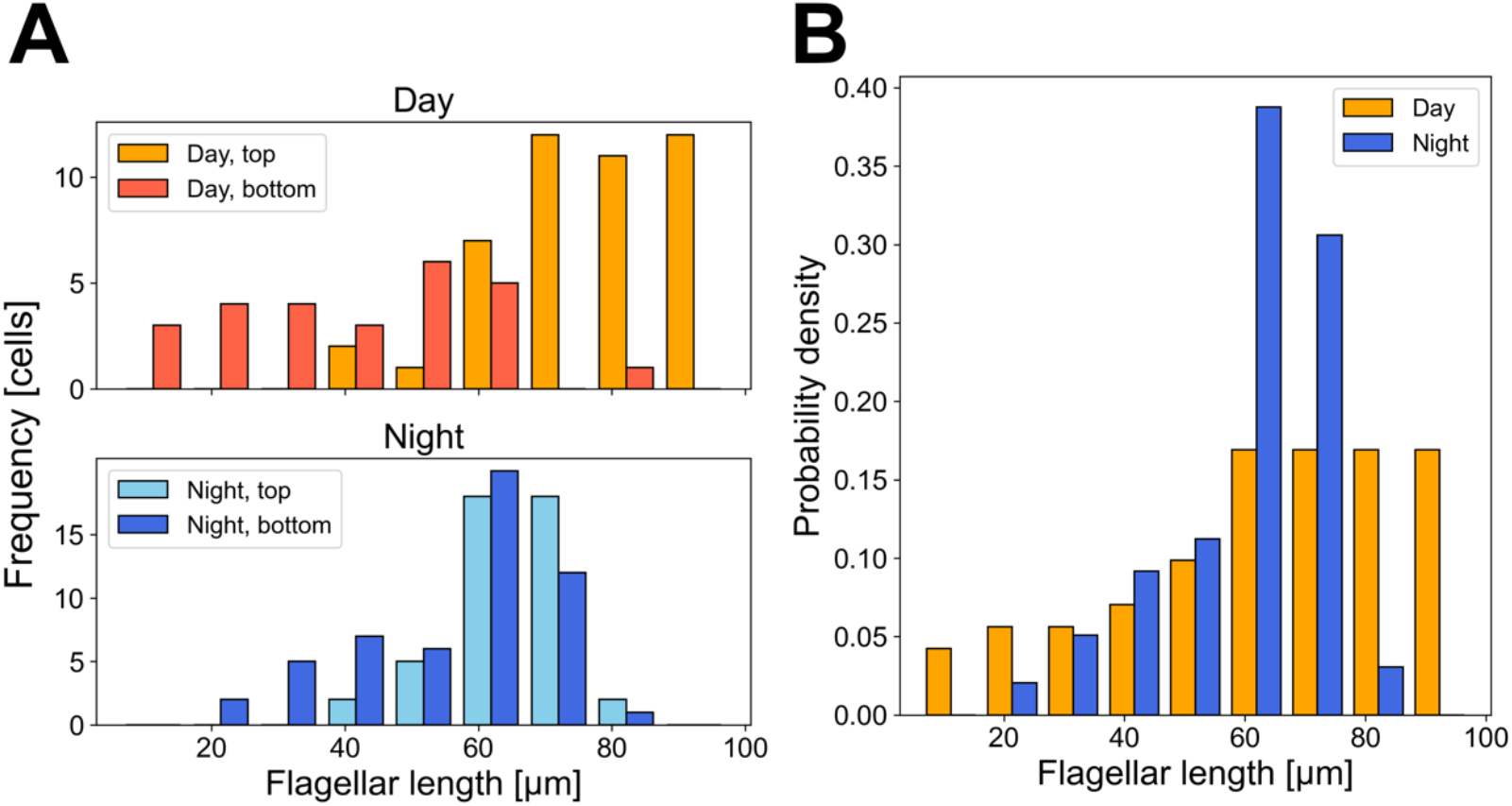
Diurnal changes in anterior flagellar length. (A) Distribution of flagellar lengths for cells observed at the top and bottom surfaces of the chamber during day and night.(B)Pooled distribution of flagellar lengths for all cells observed during the day (n=71) and night (n=98). The distributions are significantly different (Kolmogorov-Smirnov test, p=5.6×10^−4^).

A strong positive correlation was found between anterior flagellar length and swimming speed (Fig. 4). Interestingly, this correlation was stronger during the day (R=0.83, p=2.2×10^−17^, Pearson correlation coefficient) than at night (R=0.49, p=6.4×10^−7^). A Fisher’s Z-transform test confirmed this difference in correlation coefficients was significant (p=4.0×10^−5^), suggesting that at night, an additional factor (other than just flagellar length) might contribute to a reduction in swimming speed.

**Figure 4.**
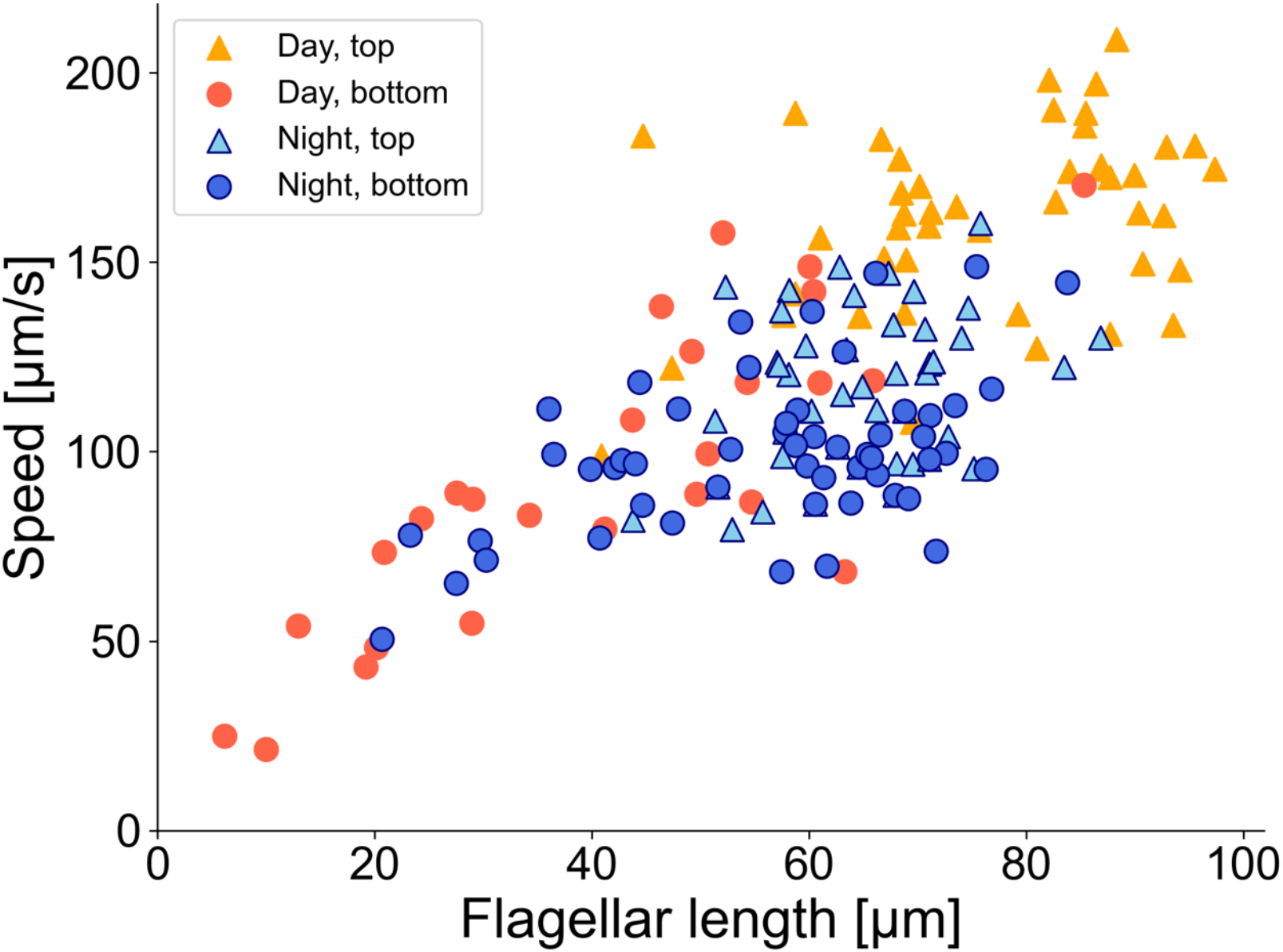
Correlation between anterior flagellar length and swimming speed. Data is shown for cells sampled during the day (n=71, R=0.83, p=2.2×10^−17^) and night (n=98, R=0.49, p=6.4×10^−7^). The correlation coefficients between day and night groups are significantly different (Fisher’s Z-transform, p=4.0×10^−5^).

### Contribution of Cell Volume

To determine if cell size influenced swimming, we calculated cell volume and compared it to swimming speed (Fig. 5). We found no evidence that larger cells swam slower due to increased drag. In fact, there was a weak positive correlation between volume and speed (day: R=0.24, night: R=0.22). Partial correlation analysis, controlling for cell volume, showed that the correlation between flagellar length and speed remained high (day: R=0.81, night: R=0.44). Conversely, the correlation between volume and speed, controlling for flagellar length, was negligible (day: R=-0.077, night: R=-0.021). These results indicate that cell volume is not a primary determinant of swimming speed in this context.

**Figure 5.**
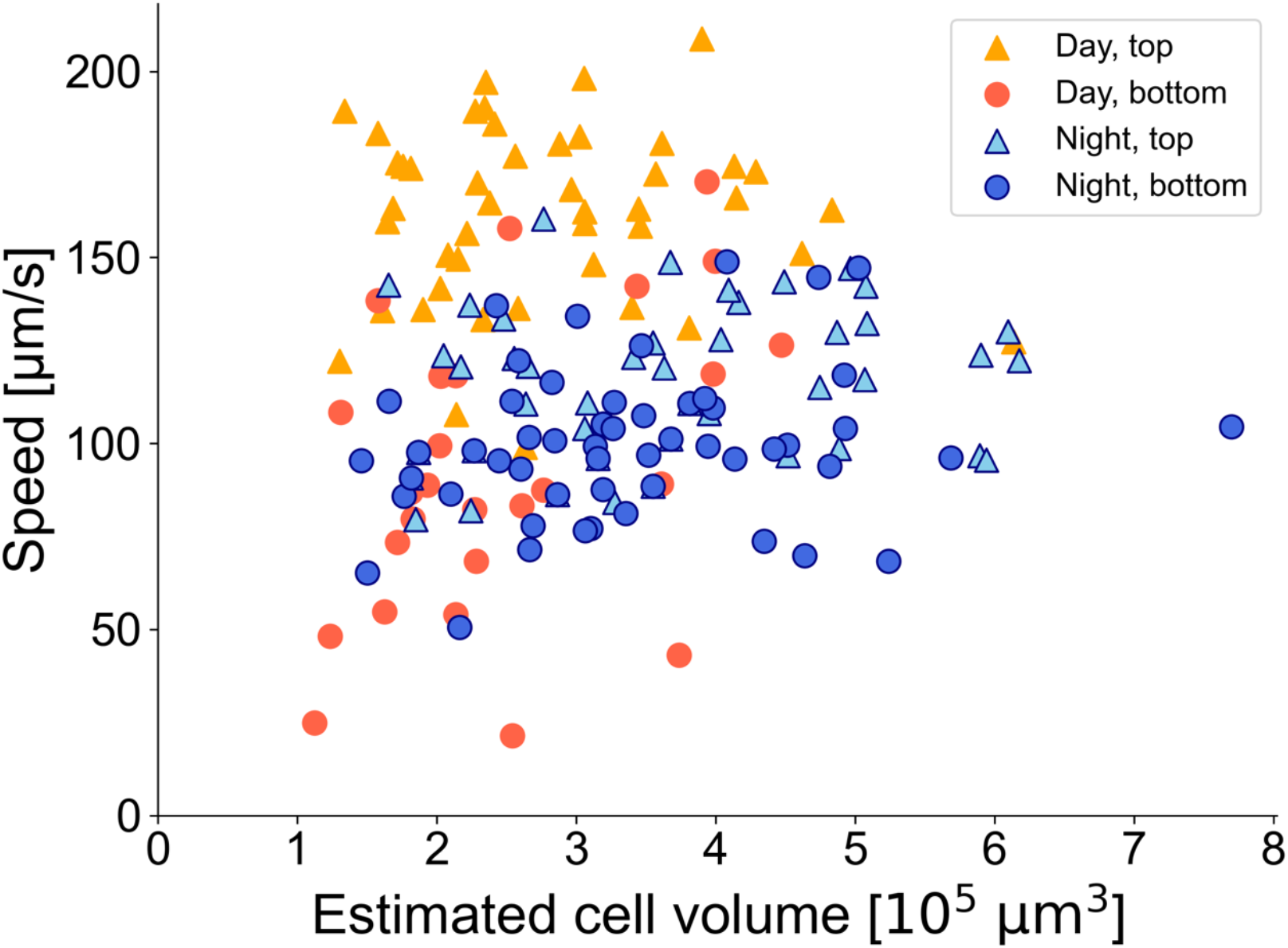
Relationship between calculated cell volume and swimming speed for day and night samples.

### Diurnal Changes in Flagellar Beat Frequency

We hypothesized the additional factor affecting night-time swimming speed might be a change in flagellar beat. We analyzed high-speed videos to determine the beat frequencies of both the anterior and posterior flagella (Fig. 6).

**Figure 6.**
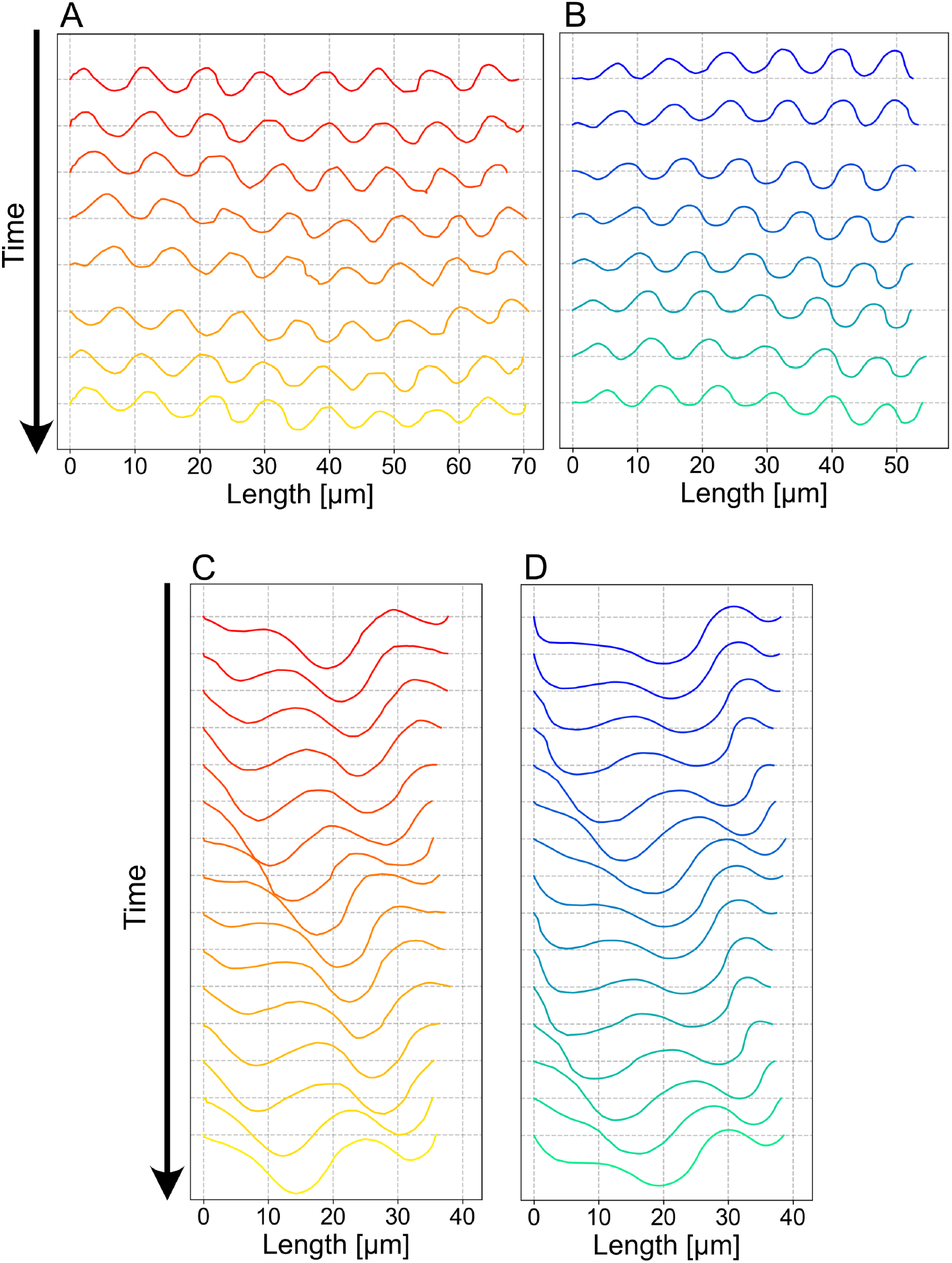
Representative flagellar waveforms. (A, B) Anterior flagellum (recorded at 200 fps) during day (A) and night (B). Traces show 8 frames (40 ms) of movement. The position of 0 µm corresponds to the base of the flagellum. (C, D) Posterior flagellum (recorded at 500 fps) during day (C) and night (D). Traces show 15 frames (30 ms) of movement. The position of 0 µm corresponds to the base of the flagellum.

The mean beat frequency of the propulsive anterior flagellum was significantly higher during the day (35.3 ± 4.9 Hz, n=11) than at night (30.6 ± 4.6 Hz, n=13) (Mann-Whitney U-test, p=0.014). In contrast, the beat frequency of the posterior flagellum showed no significant difference between day (64.6 ± 6.1 Hz, n=11) and night (62.4 ± 4.7 Hz, n=15) (p=0.41). This demonstrates that *C. marina* reduces beat frequency of the anterior flagellum at night. Although the sample size was limited, no significant correlation was found between anterior flagellar length and beat frequency (day: R=0.49, p=0.12; night: R=-0.27, p=0.36; Fig. 7).

**Figure 7.**
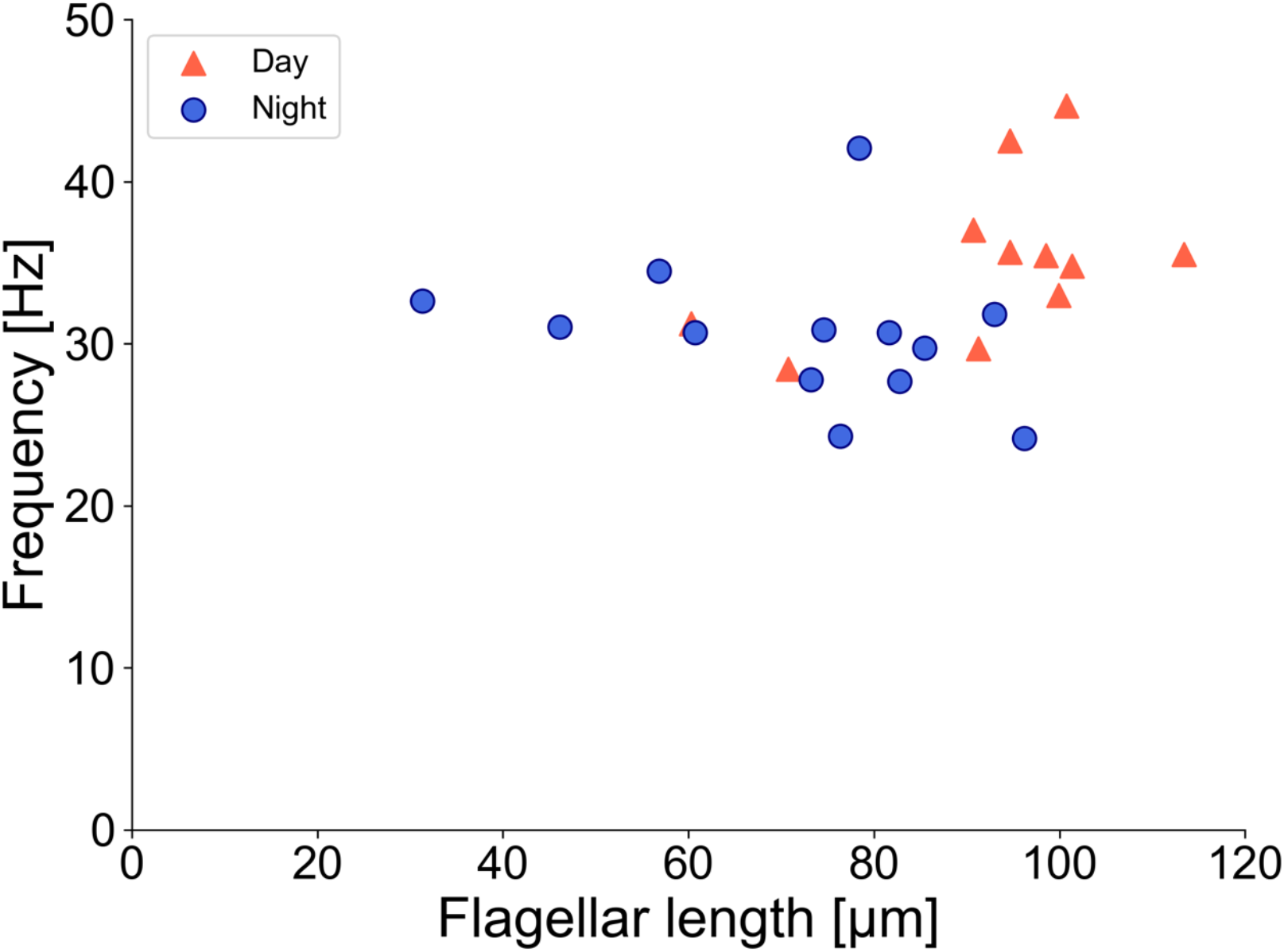
Relationship between flagellar length and beat frequency for day and night samples.

### Pharmacological Regulation of Flagellar Length

The observed diurnal change in flagellar length suggests an active regulatory mechanism. We hypothesized that this involves intraflagellar transport (IFT). To test this, we applied ciliobrevin D, an inhibitor of IFT dynein (a motor for retrograde IFT).

Treatment with ciliobrevin D resulted in a clear time- and concentration-dependent shortening of the anterior flagellum over a 6-hour period (Fig. 8). Compared to the control and DMSO-vehicle groups, which showed minimal change (7.4% in control and 14.1% in DMSO after 6 hours), the 10 µM ciliobrevin D group exhibited a mean flagellar shortening of 42.3% after 6 hours. The 5 µM group showed an intermediate shortening of 22.3%. This induced flagellar shortening was also associated with a decrease in swimming speed (Fig. 9; R=0.57, p=1.4×10^-23^, n=255).

**Figure 8.**
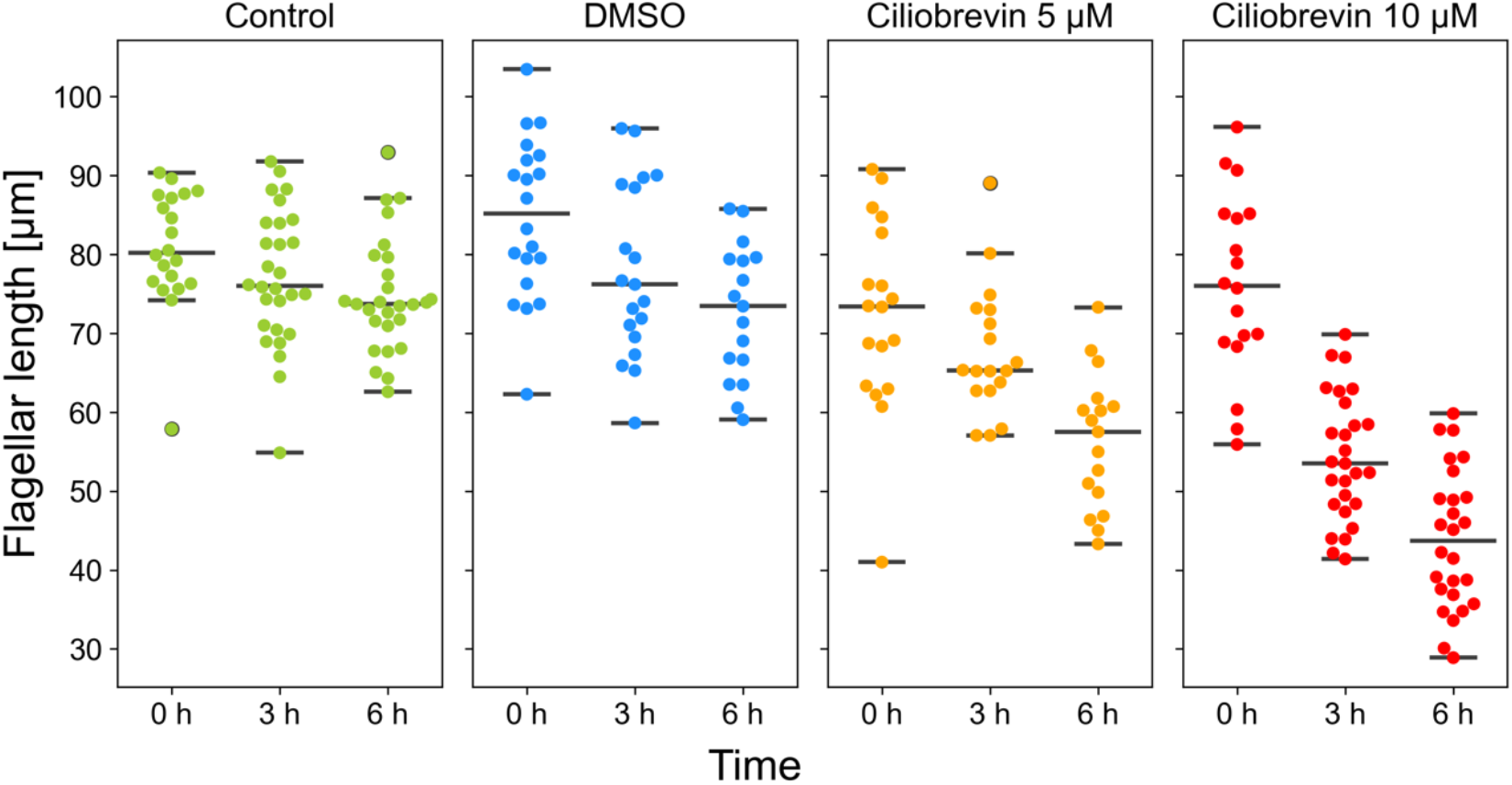
Effect of ciliobrevin D on anterior flagellar length over time. Measurements were taken at 0, 3, and 6 hours post-treatment. Groups shown: Control (no addition), DMSO (vehicle control), Ciliobrevin 5 µM, and Ciliobrevin 10 µM. Each dot represents a single cell. The horizontal lines indicate the median (middle line), along with the lowest and highest values that fall within 1.5 times the interquartile range from the first (Q1) and third (Q3) quartiles.

**Figure 9.**
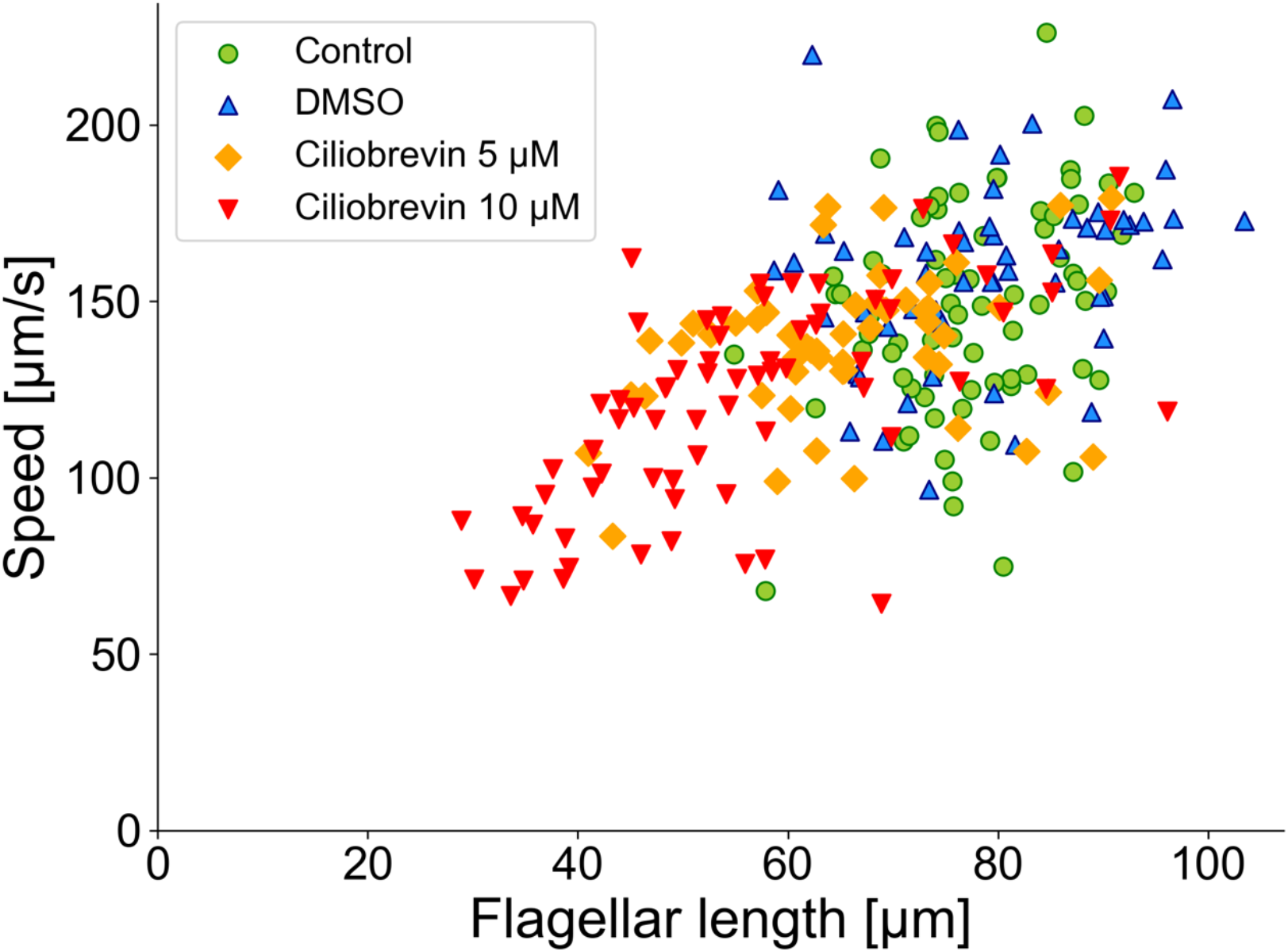
Relationship between anterior flagellar length and swimming speed in inhibitor-treated cells. Data from all time points (0, 3, and 6 hours) and all treatment groups are plotted.

## Discussion

Our study demonstrates that the DVM of *Chattonella marina* is coupled to distinct, single-cell physiological changes. We show that *C. marina* actively modulates its swimming speed by regulating two key parameters of its propulsive anterior flagellum:

1. Flagellar Length: the distribution of flagellar length was significantly different between samples during the day and at night (Fig. 4).
2. Beat Frequency: The flagellar beat frequency is significantly higher during the day and lower at night (Fig. 6a).

These two changes work in concert, resulting in faster swimming during the day, which is necessary to overcome gravity and remain in the sunlit surface layer, and slower swimming at night. This switch to a “low-power” mode (slower-beating flagella) at night is likely an energy conservation strategy. This dual-regulation mechanism also explains our finding that the correlation between flagellar length and speed was weaker at night (Fig. 4). During the day, speed appears to be primarily dictated by length (R=0.83), but at night, the significant reduction in beat frequency acts as an additional, dominant factor reducing the swimming speed, thus weakening the sole reliance on length.

Furthermore, our pharmacological experiment provides the first evidence for the mechanism underlying this length modulation. The fact that ciliobrevin D, a dynein inhibitor associated with retrograde IFT, induces flagellar shortening (Fig. 8) strongly suggests that flagellar length in this motile raphidophyte is not static but is actively maintained, likely related to IFT. While IFT is well-studied in primary cilia and the assembly of flagella (Hsiao et al., 2012; Pedersen et al., 2008), its role in the active, diurnal regulation of motile flagellar length is a novel finding.

However, our inhibitor study has limitations. Ciliobrevin may also affect other dyneins than IFT dynein, such as the axonemal dynein responsible for flagellar beating (Wada et al., 2015). Although we applied ciliobrevin concentrations that showed minimal effects to motility in sea urchin sperm (Wada et al., 2015), the observed decrease in swimming speed upon inhibitor treatment (Fig. 9) may result from a combination of flagellar shortening and a direct inhibition of the beat machinery. Further studies would be necessary to fully confirm the role of IFT in this process. Additionally, our experiments were conducted in shallow observation chambers. Future studies using taller columns that better mimic the natural environment are needed to fully understand the link between these cellular changes and large-scale vertical migration.

The present study first quantitively described the swimming characteristics of raphidophyte at the cellular scale. While the anterior flagellum thrusts the body, the role of the posterior flagellum remains to be elucidated, as there was no significant difference in the frequency of the posterior flagellum during the day and at night. In brown algae, which are multicellular organisms phylogenetically close to raphidophyte (Derelle et al., 2016), gametes have two unequal (anterior and posterior) flagella as in raphidophyte. In the male gamete of the brown alga *Mutimo*, the anterior flagellum thrust the body while the posterior flagellum involves in direction changes, with different waveform and different sensitivity to Ca^2+^ (Kinoshita-Terauchi et al., 2024). Although the present study did not capture the motion of the posterior flagellum during direction changes, the posterior flagellum of *Chattonella* might also have the role in turning. Further study should elucidate the role of the posterior flagellum in *Chattonella*.

## Conclusions

In conclusion, our findings provide a new single-cell perspective on the complex survival strategy of *C. marina*. The active, diurnal control of both flagellar length and beat frequency represents a key mechanism underlying its DVM, and our data opens a potential avenue for understanding the molecular control of flagellar length in motile eukaryotes.

## Supplementary Material Legends

**Supplementary Movie S1**. Swimming of *Chattonella marina*. Play back speed, 1/10.

## Acknowledgements

*Chattonella marina* (NIES-1) was provided by the National Institute for Environmental Studies through the National Bioresource Project (NBRP) of the Ministry of Education, Culture, Sports, Science and Technology (MEXT), Japan.

The authors thank Dr. Tomoyuki Shikata (Fisheries Technology Institute) for providing seawater and expert advice on *Chattonella* cultivation; Dr. Shoji A. Baba (Ochanomizu University) for providing the software Bohboh and Bohboh-L; Dr. Shinji Kamimura (Chuo University) for providing the stock of ciliobrevin D and invaluable advice on its experimental application, and for critical reading of the manuscript; Drs. Yuuko Wada (Ochanomizu University) and Nana Kinoshita-Terauchi (Hokkaido University) for discussions.

The authors utilized Gemini 2.5 Pro to translate and summarize the master thesis written in Japanese by Yusaku Fujita. The text was thoroughly reviewed and edited by the authors.

## Author Contributions

A.K. conceived the research. A.K. and T.N. designed the research. T.N. supervised the research. A.K. and T.N. acquired funding. Y.F. conducted the experiments. Y.F. and A.K. analyzed and visualized the data. Y.F. drafted the original Japanese text. All authors contributed to writing.

